# Hydration layer of only few molecules controls lipid mobility in biomimetic membranes

**DOI:** 10.1101/2021.05.07.443086

**Authors:** Madhurima Chattopadhyay, Emilia Krok, Hanna Orlikowska, Petra Schwille, Henri G. Franquelim, Lukasz Piatkowski

## Abstract

Self-assembly of biomembranes results from the intricate interactions between water and the lipids’ hydrophilic head groups. Therefore, the lipid-water interplay strongly contributes to modulating membranes architecture, lipid diffusion, and chemical activity. Here, we introduce a new method of obtaining dehydrated, phase-separated, supported lipid bilayers (SLBs) solely by controlling the decrease of their environment’s relative humidity. This facilitates the study of the structure and dynamics of SLBs over a wide range of hydration states. We show that the lipid domain structure of phase-separated SLBs is largely insensitive to the presence of the hydration layer. In stark contrast, lipid mobility is drastically affected by dehydration, showing a 6-fold decrease in lateral diffusion. At the same time, the diffusion activation energy increases approximately twofold for the dehydrated membrane. The obtained results, correlated with the hydration structure of a lipid molecule, revealed that about 6-7 water molecules directly hydrating the phosphocholine moiety play a pivotal role in modulating lipid diffusion. These findings could provide deeper insights into the fundamental reactions where local dehydration occurs, for instance during cell-cell fusion, and help us better understand the survivability of anhydrobiotic organisms. Finally, the strong dependence of lipid mobility on the number of hydrating water molecules opens up an application potential for SLBs as very precise, nanoscale hydration sensors.

## Introduction

Biological cell membranes are dynamic barriers composed of a large variety of lipids and embedded with various proteins. Due to the complex miscellaneous molecular interactions occurring in cellular membranes, lipid model systems are particularly attractive alternatives for the controlled investigation of various physicochemical processes affecting the membrane architecture and dynamical properties. In this regard, self-assembling supported lipid bilayers (SLBs) have been well accepted as one of the most suitable model membrane systems, due to their analogous physical and structural properties to those of biomembranes, and their easy preparation and handling methods^1^. Consequently, SLBs have been exploited to investigate membrane architecture and properties such as domain formation^2,3^, lateral diffusion or ion transport^4,5^, and biological processes at the cellular and molecular levels, such as protein-membrane interactions^6^, ligand-receptor interactions, cellular signalling^7,8^ or cell adhesion^9,10^.

Various types of chemical and physical interactions determine the complex properties, architecture, and activity of the membrane. Membrane intricacy results not only from the interactions between membrane constituents, such as lipid-lipid and lipid-protein interactions but also from the hydrophobic mismatch that arises from the interplay with water hydrating the membrane. In fact, hydrophobic mismatch is considered to be one of the key physicochemical mechanisms that regulate membrane organization and promotes nanoscopic and microscopic separation of liquid ordered (L_o_) and liquid disordered (L_d_) phases, and determines the position and orientation of transmembrane proteins in both model and living cell membranes^6^. The thin layer of water that directly hydrates the membrane, commonly referred to as biological water^11^, has been proven to actively participate in the biological functioning of proteins and DNA^12^. Numerous experiments aimed at understanding the properties of biological water, using nuclear magnetic resonance^13^, X-ray and neutron scattering^14,15^, infrared spectroscopy^16^, sum frequency generation^17,18^ and molecular dynamics simulations^19–22^, among others, showed that water molecules form a network structure, being bound by hydrogen bonds and weak van der Waals interactions around the polar headgroup – the so-called clathrate hydration structure^19^. Moreover, water molecules present in the direct or indirect hydration shell around the head group region exhibit markedly different properties from those of bulk water^23^.

Water is unambiguously essential for maintaining biological activities in living systems. But, in fact, nature shows various phenomena of anhydrobiosis: ‘life without water’ in which the cell membrane not only survives harsh dehydration but also regains full activity upon rehydration. The most common method allowing dehydration is an increased production of carbohydrates (mostly trehalose) in organisms such as tardigrades^24–27^, nematodes^28^, and yeasts^29,30^. The water-replacement hypothesis states that trehalose stabilizes the head groups and enables maintaining the spacing between the fatty acyl chains^31^. On the other hand, *Bdelloid rotifers* base their survival mechanism on the contraction of the body, which reduces the surface exposed to the environment and allows slow evaporation^32,33^. High desiccation resistance in seeds, pollens and anhydrobiotic plants is associated with the production of LEA (late embryogenesis abundant) proteins that are responsible for ion sequestration, protection of membrane, and renaturation of proteins that unfolded due to the lack of water^34^.

Importantly, it should be noted that local, temporary dehydration of the cell membranes also occurs continuously in our bodies during, e.g., adsorption of biomacromolecules or cell-cell fusion events. A prerequisite for membrane fusion is establishing close contact between the outer leaflets of lipid bilayers such that the thin layer of water molecules is expelled (the clathrate hydration structure is disturbed) and finally overcoming the energy barrier, commonly referred to as ‘hydration force’, present mainly due to the repulsive forces between lipid bilayers^35,36^.

Last but not least, various studies of the electrical, mechanical and physicochemical properties of planar lipid bilayers have revealed that these platforms have huge application potential from a technological standpoint as biosensors and bio-coatings^37,38^. These bio-applications require SLBs to be exposed to changes of external conditions such as temperature during preservation, reagent addition, and, importantly, humidity.

Hence, understanding the interplay between the hydration layers and the cell membrane is of utmost importance, both in unraveling mechanisms behind membrane organization and activity and in the frameworks of biotechnology and bioengineering. Unfortunately, the ability to investigate the intimate interactions between the membrane and the biological water has so far been hindered by the lack of appropriate experimental approaches for the preparation and study of lipid membranes in a controlled hydration state. In particular, keeping the membrane structure intact under decreased hydration conditions is challenging^19,21^. So far, several approaches to protect the membrane from rupturing and vesiculation have been utilized: modification of lipid head groups in order to strengthen the SLB-mica attractive interactions^39–42^, cross-linking the lipid bilayer^43,44^, attaching polymers to the head group of lipids^39,45^ and adding biomolecules like proteins, disaccharides or enzymes^38,44,46–48^. These approaches, however, inevitably alter the intrinsic properties of the SLBs. Moreover, the exact hydration state of the membrane is unknown. Consequently, a method for preparing and stabilizing the membrane under varying, well-controlled hydration conditions without the use of additional stabilizing agents or chemical modification is needed.

Here, we present an unprecedented way to obtain phase-separated, stable SLBs with a well-controlled hydration state without interfering with membrane composition, which enables the investigation of bilayer structure and dynamics under arbitrary hydration conditions. Using a combination of fluorescence microscopy imaging and fluorescence recovery after photobleaching (FRAP) experiments, we report interesting observations of the structural and dynamical changes taking place. We show that the structure of SLB can be preserved under dry conditions by a controlled drying process with a slow and sequential reduction in relative humidity of the membrane environment. Such an approach revealed that the lateral diffusion dynamics of the liquid disordered phase is significantly reduced with dehydration. Importantly, the membrane can undergo multiple de- and rehydration cycles always reviving its native dynamics. We also show that the diffusion activation energy for lipids in a dehydrated membrane is much higher than for fully hydrated SLB. Finally, we provide molecular-level insights into how and which water molecules around lipids play a key role in regulating lipid dynamics in the membrane.

## Experimental Section

### Materials

1,2-dimyristoleoyl-*sn*-glycero-3-phosphocholine (14:1 PC), egg yolk sphingomyelin (SM), 23-(dipyrrometheneboron difluoride)-24-norcholesterol (TopFluor cholesterol) were purchased from Avanti Polar Lipids, Alabaster AL., USA. Monosialoganglioside (GM1) from bovine brain and 1,2-dioleoyl-*sn*-glycero-3-phosphoethanolamine labeled with Atto 6_33_ (DOPE-Atto 6_33_), 4-(2-hydroxyethyl)piperazine-1-ethanesulphonic acid (HEPES) sodium salt, and sodium chloride (NaCl) were purchased from Merck KGaA, Darmstadt, Germany. Alexa Fluor 488 conjugated with cholera toxin B subunit (CTxB 488), Alexa Fluor 594 conjugated with cholera toxin B subunit (CTxB 594) were obtained from Molecular Probes, Life Technologies, Grand Island, NY, USA. All the materials were used without further purification.

### Vesicle preparation

The SLB was prepared by vesicle deposition method following formerly established protocol^49^ with suitable modification. In order to form multilamellar vesicles (MLVs), 14:1 PC, SM and cholesterol in chloroform solution were mixed at a molar ratio 1:1:1 with the addition of 0.1 mol% of GM1 and 0.1 mol% of DOPE-Atto-6_33_ to form 10 mM solution of the lipids. The lipid mixture was dried under nitrogen gas leaving a thin film of lipids deposited on the bottom of the vial, followed by desiccation under vacuum for at least 2 hours. The lipids were resuspended in buffer solution (10 mM HEPES and 150 mM NaCl, pH adjusted to 7.4) and exposed to few cycles of heating on the hot plate at 60°C and vortexing. Lipid suspension containing MLVs was aliquoted into sterilized glass vials and diluted 10 times (final concentration of lipids 1 mM) using buffer solution. Aliquots were stored at −20°C for further use.

### SLBs preparation

Aliquots containing MLVs of the desired composition were bath-sonicated for 10 min at maximum power to generate small unilamellar vesicles (SUVs). Freshly cleaved mica was glued to a coverslip by UV-activated glue (Norland 68) and the top layer of mica was removed right before the deposition to keep the surface properties of freshly cleaved mica intact during deposition. A half-cut Eppendorf tube was placed on the top of the coverslip and sealed with silicone. 100 μL of SUVs solution was deposited on top of mica followed by the addition of 2 μL of 0.1 M CaCl_2_ solution and 9 μL of 0.01 mM CTxB dissolved in buffer solution, all at room temperature. The SLB was allowed to settle for 30 sec, and then 400 μL of buffer solution (10 mM HEPES and 150 mM NaCl, pH adjusted to 7.4) was added and the sample was incubated for 30 minutes. The bilayer was rinsed 10 times with 2 mL of buffer solution to wash out excess unfused vesicles. The Eppendorf tube reservoir was fully filled with buffer solution, closed with a glass coverslip and sealed with silicone to prepare a fully hydrated sample containing bulk water.

### Preparation of SLBs at different hydration level

In our work, two distinct methods were implemented for drying the SLBs. (a) The bulk water was pipetted out and the sample was left open to dry and equilibrate to atmospheric humidity (~30% RH) at room temperature and (b) after removal of bulk water by micropipette the sample was equilibrated in atmosphere of different relative humidity (RH%). Atmosphere of different relative humidity was created inside the open half-cut Eppendorf tube by purging nitrogen gas of specific relative humidity using a homebuilt control unit (see Fig. 1A). The setup consisted of three flowmeters, three manual valves, a reservoir with water and an electronic hygrometer with 0-95% RH range and 1% precision. Relative humidity of nitrogen gas was adjusted and maintained by mixing a suitable amount of wet (saturated with water vapor, 90% RH) N_2_ and dry (2-3% RH) N_2_ gas. The electronic hygrometer was used to monitor the final relative humidity and temperature of the N_2_ gas being purged towards the sample. To study SLB structure and dynamics at different relative humidity, the silicone seal of the sample was cut and water was pipetted out completely and purging of wet nitrogen gas of 90% RH was started immediately towards the SLB. RH was decreased (and subsequently increased) in steps of ~20% at a rate of 2-3% RH per minute. Next, the SLB was equilibrated at a given RH for about 10 minutes before FRAP measurements were performed. Relative humidity of wet nitrogen gas was decreased gradually from 90% (62·10^19^ water molecules/min) to approximately 70% (48·10^19^ water molecules/min), 50% (34·10^19^ water molecules/min), 30% (20·10^19^ water molecules/min), and finally dry nitrogen (around 2-3% RH) was purged to the SLB. Similarly, rehydration of the dried SLB was done by purging wet nitrogen gas with increasing relative humidity and finally resealing the half-cut Eppendorf tube filled with water.

**Figure 1.**
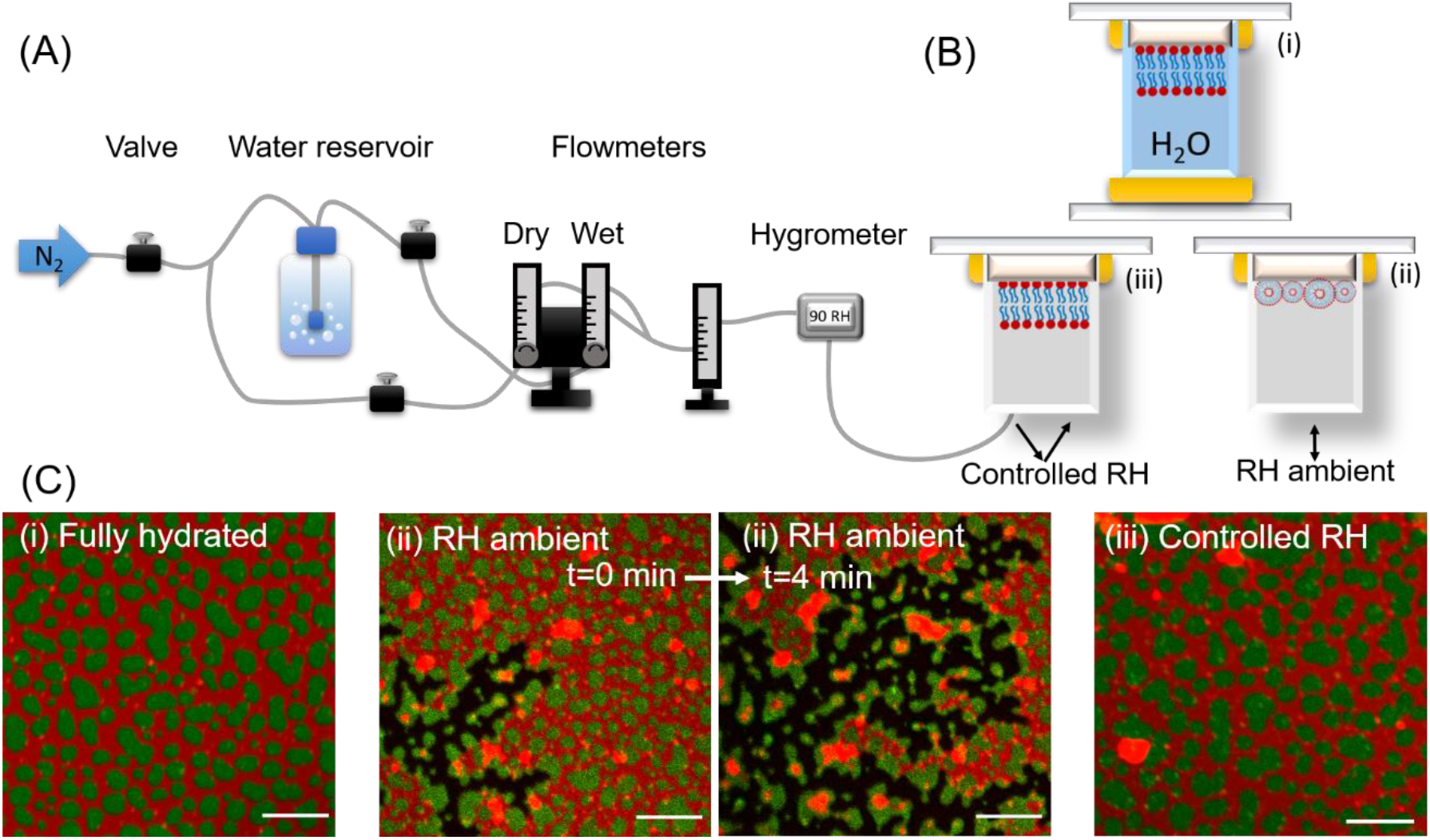
Schematic representation of the home-built humidity-controlling set-up. (B) Cartoon depiction of the three types of SLBs studied here: (i) fully hydrated with bulk water, (ii) exposed to the ambient humidity, (iii) exposed to atmosphere with well controlled humidity. (C) Fluorescence images of the representative SLBs in different hydration conditions indicated in panel B. The two middle panels show progressive rupturing and delamination of the SLB abruptly exposed to ambient RH. Scale bar corresponds to 10 μm.

### Fluorescence Microscopy and FRAP

Laser-scanning confocal imaging and FRAP experiments were performed on SLBs using upright Zeiss LSM 710 (Carl Zeiss, Jena, Germany) microscope with 40× 1.3 NA oil immersion objective. Lasers of wavelengths 488 nm and 633 nm were used for excitation of Alexa Fluor 488 and Atto-633-DOPE respectively. In the case of 3-fold labeling with TopFluor cholesterol, CTxB-Alexa Fluor-594 and Atto-633-DOPE, lasers of 488 nm, 543 nm and 633 nm were applied accordingly. Laser power was adjusted during imaging to avoid excessive photobleaching of the sample. Small circular spot of 10 μm diameter was bleached and the area of the bleached spots was kept constant for all FRAP experiments. Diffusion coefficients were calculated by fitting the fluorescence recovery curve considering free Brownian lateral diffusion of lipid molecules in the membrane using modified Soumpasis formula^50^: *F(t)* = *a + b* · *f(t)*, where *a* is the amplitude of the recovery function, *b* is the remaining fluorescence after bleaching and *f(t)* is the Soumpasis function. Fitting was done for data normalized with respect to the reference intensity signal of the whole image excluding the bleached spot. Complete dehydration and rehydration cycle was performed for 3 samples and FRAP experiments were performed in at least 5 different spots at a particular relative humidity for each sample. Diffusion coefficients were averaged over a range of RH at which particular traces were measured.

### Temperature dependence experiments

Variation of D(T) was examined for two hydrated and two dehydrated samples in the temperature range 25±1 - 45±1 °C. A resistive tape was attached to the sample reservoir tube for heating and a thermocouple has been placed inside the reservoir tube for continuous monitoring of sample temperature. Activation energies for the two samples were calculated using Arrhenius equation: *lnD = lnA − E_α_/RT* where D is the diffusion coefficient, A is the pre-exponential factor (assumed to be temperature independent in this range), R is the universal gas constant and T is the temperature in Kelvin scale. For Arrhenius plots, weighted linear regression of lnD values was presented. The confidence bounds generated by the fitting of FRAP traces were considered as error bars for D and their reciprocals to be the weights.

## Results

### Structure

In this study, the changes in the membrane structure at different hydration conditions have been examined by fluorescence imaging. In the experiments we considered three levels of membrane hydration, described in detail in the experimental section and schematically depicted in Fig. 1B: (i) fully hydrated SLB, where the membrane is submerged in bulk water, (ii) SLB for which most of the bulk water was pipetted out and the sample was left open to equilibrate to room humidity (~30% RH) and (iii) SLB for which bulk water was removed to the highest extent and the sample was immediately exposed to N_2_ atmosphere with ~90% RH.

Fully hydrated SLBs, marked as (i) in Fig. 1B–C, exhibit homogeneously distributed domains of liquid-ordered (L_o_) phase with an average area of 1.77 ± 0.29 μm^2^. The domain size, distribution, and shape are typical to L_d_/L_o_ phase-separated SLBs prepared in such conditions, in full agreement with previous reports^51^. SLBs are far from static – over time they slowly merge with each other to form bigger domains. Over the course of ~24 h the average domain size increases by up to 40%.

The SLB (marked as (ii) in Fig. 1B–C) for which most of the bulk water was removed and the surface was exposed to open air of low RH (~30%) initially exhibits identical structure to the fully hydrated sample. Compared to fully hydrated SLBs, here we observed an increase in vesicle-like aggregated structures, mainly composed of L_d_ phase residing on the surface of the SLB. However, as spontaneous drying proceeds, the remnant bulk water layer shrinks, causing the drop-like macroscopic water layer wavefront to pass over the membrane surface. The local changes of surface tension induce delamination of the membrane from the mica support (the two middle panels in Fig. 1C). Intriguingly, in most cases, L_d_ phase detaches from mica first, while L_o_ domains remain attached to mica (extended time series is shown in Fig. S1 and in movie M1). Shortly after, over the course of a few minutes, also L_o_ domains shrink up and form curled-up vesicle-like structures mixed up with the L_d_ phase lipids. Delamination of the membrane ceases as soon as the residual bulk water is evaporated. However, it should be noted that in several areas confined by mica terraces, SLB structure remains unperturbed (Fig. S2).

Markedly different behavior was observed when the SLB was exposed to N_2_ atmosphere with a high RH of ~90%, directly after bulk water removal. SLB kept under a continuous flow of N_2_ atmosphere with high RH (denoted as (iii) in Fig. 1B–C) qualitatively closely resembles fully hydrated SLB. Minor delamination was observed solely on the perimeter of the sample, close to the mica edges. This curling up of the membrane occurs during the time required to remove bulk water and expose the membrane to an atmosphere of high RH. These events are likely responsible for the increased number of vesicles and aggregates at the top of the membrane (Fig. 2A–2C). The aggregates that appear due to bulk water removal are initially mobile and float while the residual water evaporates. Once the sample equilibrates with an atmosphere of high RH (70-80% RH), the aggregates become stagnant. No change in the structure or quality of the SLB kept in such conditions was observed over the course of a few hours. No significant change in the quality of the membrane structure was noticed, although the perimeter of domains became increasingly jagged with further, gradual decrease of RH down to about 50%.

**Figure 2.**
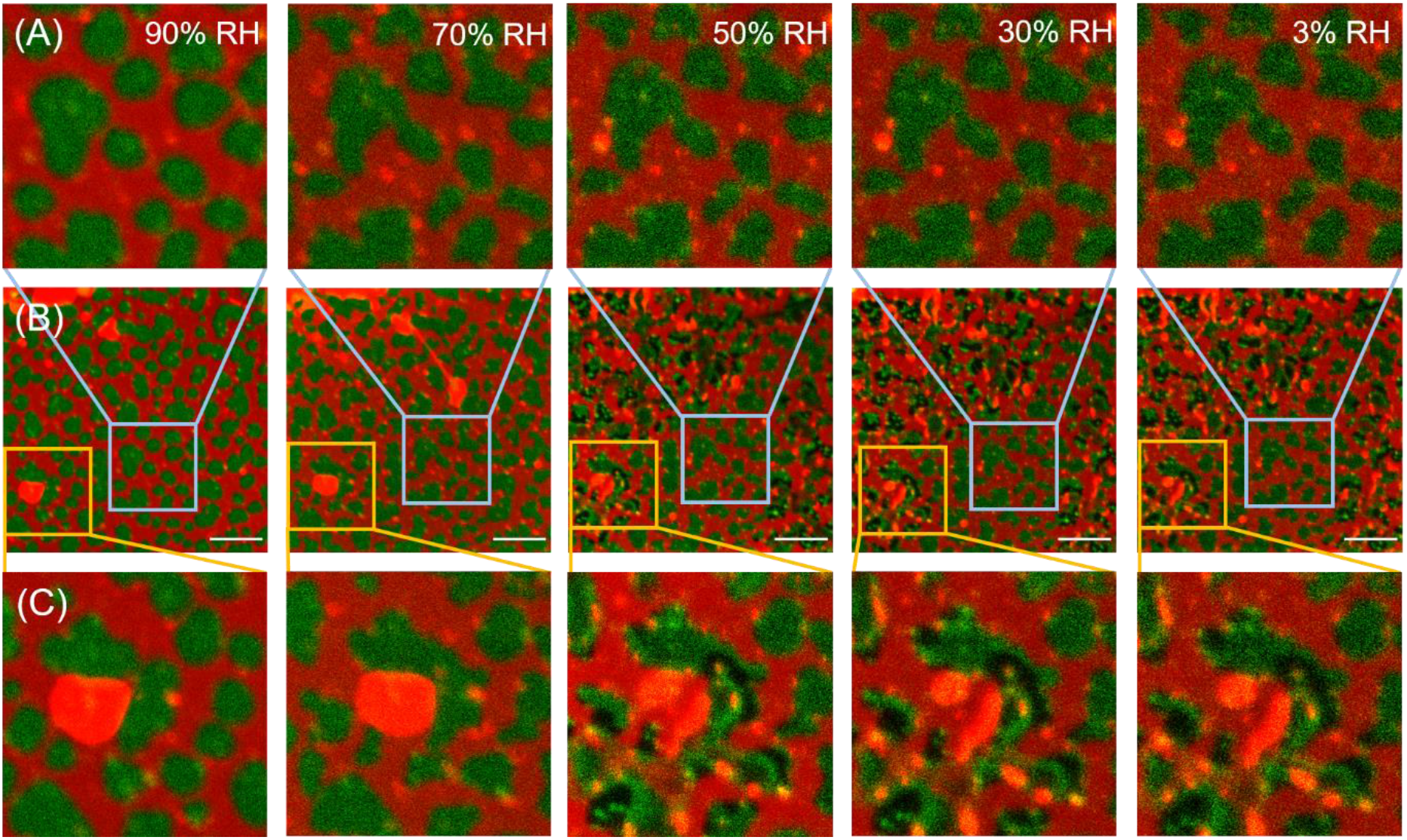
Consecutive fluorescence images of the same area of SLB exposed to 90%, 70%, 50%, 30%, and 3% RH. Top row (A) and bottom row (C) show the zoomed-in region indicated by the blue and yellow rectangles in images in the middle row (B), respectively. Equilibration time for each hydration condition and between consecutive images was ~30 minutes. Scale bar corresponds to 10 μm.

At around 50% relative humidity, the appearance of seemingly hole-like dark spots within L_o_ domains (labeled with CTxB-Alexa 488) is observed at several locations on confocal microscopy images. In the range of 50% through 30% to nearly 0% RH, the membrane structure does not change significantly except the appearance of the dark spots in L_o_ domains in few more locations. Noticeably, the formation of these hole-like dark spots is limited to few areas, while an unperturbed and continuous phase-separated membrane structure can be observed over the prevalent sample area even at relative humidity close to 0%. Evidently, by means of slow, well-controlled and gradual (~2-3% RH/min – for details see Experimental section) decrease of membrane hydration, air-stable membrane can be formed without the addition of external stabilizing agents. Additional confocal images of the sample as a function of hydration are shown in supplementary information (Fig. S3). With lowering hydration below 50% RH the big aggregates (ranging from 5-25 μm^2^), located at the top of the membrane, break into smaller ones.

It should be noted that the membrane equilibrated at different hydration states is stable for up to a few hours. Intriguingly, the process of dehydration is fully reversible, i.e. dehydrated membrane can be rehydrated back to the state compliant with high RH and further to full hydration by addition of bulk water. Upon rehydration, the darker spots within L_o_ domains become homogeneously bright again and the domains regain their former (rounder) shapes at around 70-85% RH. Images of SLB at different RH during rehydration are shown in Fig. S3.

### Dynamics

Next, we examined whether the hydration state of the membrane affects the mobility of the lipids by performing FRAP experiments on membranes equilibrated at different hydration conditions. The mobility of lipids constituting the membrane depends on the composition of the SLB^52^. The measured single component fully hydrated membrane of 14:1 PC shows higher diffusion coefficient (2.93 ± 0.44 μm^2^/s) than the L_d_ phase of our, phase-separated SLB (~1.7 μm^2^/s), which is consistent with the previous reports^53^.

FRAP traces obtained for the phase-separated SLBs in different hydration states are shown in Fig. 3A. Evidently, with lowering hydration of the membrane, the mobility of the L_d_ phase decreases significantly. At fully hydrated condition, i.e. before removal of bulk buffer solution, the L_d_ phase lipids showed the highest mobility of 1.66 ± 0.22 μm^2^/s. After the withdrawal of bulk water and being equilibrated to ~90% RH, the mobility remained unaltered. With a further decrease in hydration level, the diffusion coefficient (D) of lipids has been observed to decrease prominently (Fig. 3B). The average D decreases over 6 times during dehydration from 1.69 ± 0.29 μm^2^/s, for 87% ± 2% RH to 0.27 ± 0.29 μm^2^/s at 3% ± 2% RH. A steady decrease in mobility of lipids is observed from full hydration to around 50% RH. Below 50% RH, the mobility of L_d_ lipids remains almost constant. The fluorescence recovery for fully hydrated membranes and membranes equilibrated with high RH% (~90%) reaches 93% ± 3% of the initial fluorescence intensity. At relative humidity less than 85% the fluorescence does not recover up to the initial intensity and in the case of RH lower than 50%, fluorescence recovery is significantly lower and amounts to less than 50% of the initial fluorescence intensity. The extracted mobile fraction, defined as the amplitude of the fitted recovery function normalized to the total bleach depth 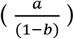, as a function of (de)hydration state of the membrane, is shown in Fig. 3C.

**Figure 3.**
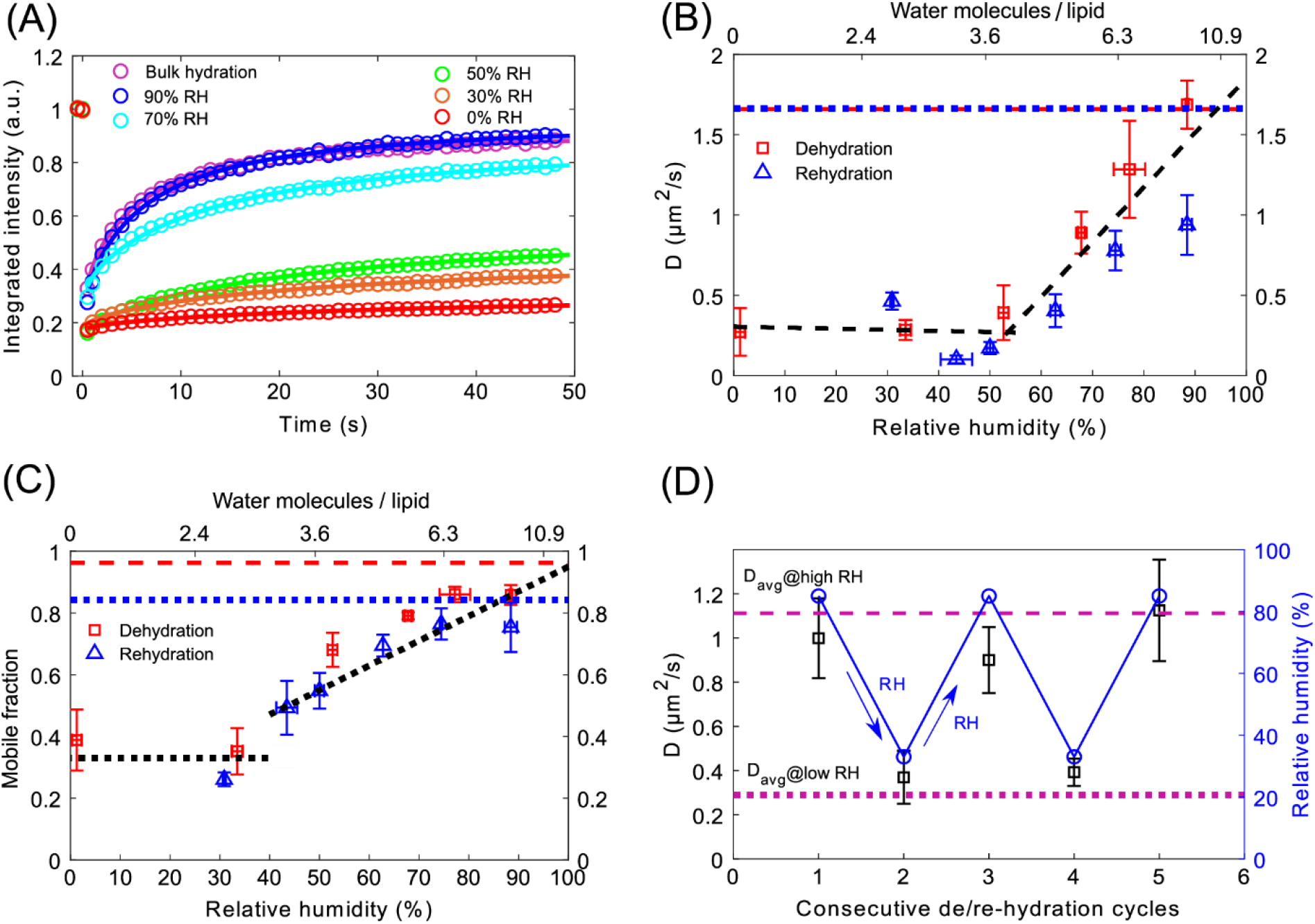
(A) FRAP traces of fully hydrated SLB and SLB equilibrated to 90%, 70%, 50%, 30% and 0% relative humidity. (B) Diffusion coefficient for the L_d_ phase for SLBs at different relative humidity during dehydration (red squares) and rehydration (blue triangles). The data points correspond to the diffusion coefficient averaged from at least 5 FRAP traces from each of the 3 samples at a particular RH. The two black dashed lines are separate linear regressions of the data points >55% RH and <55% RH. The red dashed and blue dotted lines correspond to diffusion coefficient for fully (bulk) hydrated sample before dehydration and after rehydration, respectively. (C) Mobile fractions extracted from the fits of the modified Soumpasis formula (see Methods) to the FRAP traces during dehydration (red squares) and rehydration (blue triangles). The black dotted lines are guides to the eye highlighting the data changes similar to those in panel B. The red dashed and blue dotted lines correspond to mobile fractions for fully hydrated sample before dehydration and after rehydration, respectively. (D) Diffusion coefficient (black squares) averaged over 5-7 FRAP traces at each hydration level during consecutive dehydration and rehydration (87% ↔ 33% RH) cycles (blue circles). Purple dashed and dotted lines correspond to the average diffusion coefficient for all the measured FRAP traces for SLBs kept at high (85-90% RH) and low (30-35% RH) relative humidity, respectively.

Interestingly, during rehydration of the SLB, by increasing the relative humidity level gradually from 0% to 90%, the mobility of lipids increased accordingly and was strongly correlated with the diffusion coefficient observed during dehydration of the membrane (Fig 3B).

The extracted mobile fraction during the rehydration process also closely resembles that observed during dehydration for each specific hydration state. Upon a full dehydration/rehydration cycle, both the average D value and mobile fraction reach back their initial values. Taking all the data into account, we observed two regimes. In the range of 50-90% RH, D exhibits significant changes with hydration. On the other hand, below 50% RH, D is nearly independent of the hydration of the membrane. The linear regressions performed on the data points in these two ranges show a clear turnover point at about 50% RH. A similar trend is observed for the mobile fraction – a significant decrease above 50% RH and little dependence in the hydration range below 50% RH.

Consecutive cycles of drying and rehydrating the SLB in the range of 87% to 33% RH were performed three times on the same sample while at each hydration state recording FRAP traces from a minimum of 6 spots. The sample was equilibrated for 10 minutes at a particular RH%. Remarkably, once bulk water is completely removed, the membrane exhibits very good stability in terms of structure and full reversibility of its dynamics. Keeping the membrane in such conditions allows strong modulation of the mobility by a factor of nearly 4 - ~0.3 μm^2^/s vs ~1.2 μm^2^/s (see Fig. 3D).

In accordance with previous reports the diffusion rates of L_o_ and L_d_ phases are significantly different: (1.66 ±0.22) μm^2^/s for L_d_ and (0.1 ±0.01) μm^2^/s for Lo. While qualitatively it appears that the diffusion coefficient decreases for the L_o_ phase when lowering membranes hydration, it is very difficult to quantify this change in a reliable manner for two reasons: a) the diffusion coefficient is already very low at full hydration and b) the signal to noise ratio of the signal is quite low due to much lower fluorescence quantum efficiency of the CTxB label at low hydration conditions.

## Discussion

### Structure

The multicomponent SLBs composed of 14:1 PC, SM, and cholesterol exhibit substantial structural changes with abrupt dehydration, whereas remain largely intact at lower hydration conditions when subjected to a well-controlled, gradual decrease in hydration level.

After bulk dehydration, the membrane is covered with a remnant, thin layer of water that desorbs over time. Exposing the membrane to the ambient RH causes the residual water to evaporate rapidly, causing fast shrinking of the water layer and inducing delamination and curling up of the membrane followed by lipid aggregation (see Fig. 1C, S1 and movie M1). This is due to the domination of the air-water interfacial force over the attractive forces between the mica substrate and the proximal leaflet of the SLB^38^. Detachment and curling up of the L_d_ phase prior to the L_o_ phase during drying can be explained by differences in mechanical properties of the two phases. The L_o_ phase is stiffer (higher bending modulus and area expansion modulus) than the L_d_ phase^54^, which results in lower steric forces and stronger interaction with the substrate. The observed stronger interaction of the L_o_ phase with mica than the L_d_ phase is consistent with the stronger adhesive interactions observed for DSPC gel phase domains and reported in the previous research^55^. It should be noted that the probability of survival of SLBs during rapid drying is increased by the presence of intrinsic defects of the support, such as mica terraces and/or cleaving defects (see Fig. S2). The defects obstruct the drying water, decreasing the local water-air tension and protecting the membrane from delamination. This observation is in accordance with the previous report on preparation of air-stable membrane by generating an obstacle network made of peripheral enzyme phospholipase A_2_ as physical confinement, where the presence of defects affects local surface tension and stop the water-air from propagation, leaving the membrane intact^38^.

In contrast, for the SLB exposed and equilibrated to high relative humidity (~90%) the overall membrane structure remains largely unaffected, except for the deposition of a few aggregates on top of the bilayer (see Fig. 2). Upon decreasing the relative humidity further in the range of 90-55% we observed no significant changes to the structure of the membrane – the SLB still exhibits homogeneously distributed L_o_ domains in a L_d_ matrix. With decreasing hydration, however, the perimeter of the L_o_ domains becomes increasingly jagged (see Fig. 2A). The ragged outlines of the L_o_ domains are mostly evident in the AFM topography image acquired on a membrane equilibrated to 30% RH (see Fig. S5). AFM studies of fully hydrated SLBs of analogous composition showed round L_o_ domains with smooth perimeter^56^. Moreover, the thickness difference between the L_d_ and L_o_ phases for dehydrated SLB is nearly 3 times lower (~0.6 nm, see Fig. S5) compared to the thickness mismatch for fully hydrated SLB with the same composition (~1.56 ± 0.13 nm)^56^. Clearly, lowering the hydration of the membrane leads to a decrease in the hydrophobic mismatch between the L_d_ and L_o_ phases, and consequently of the line tension.

At lower hydration conditions (<50% RH), dark spots in some of the L_o_ domains appear (see Fig. 2C), where fluorescence of the labeled GM1-CTxB complex is not detected. At the same time, parts of these L_o_ domains exhibit locally higher fluorescence intensity. Detailed analysis of the fluorescence images reveals the nature of the dark spots within the L_o_ domains. The shape (outline) of the domains before the appearance of the dark spots (RH>50%), with the dark spots present (RH<50%), and after the disappearance of the dark spots (upon rehydration) remains the same (see Fig. S6). If the dark spots were due to the formation of holes within the membrane, one would expect that upon rehydration the shape would randomly change, i.e., the holes would be filled randomly by L_d_ and/or L_o_ phase. Instead, we observe that the L_o_ domains maintain their original shape and regain fluorescence distribution as before the dehydration.

Next, we analysed the fluorescence intensity of selected L_o_ domains containing the dark spots as a function of hydration. The total integrated fluorescence intensity of an L_o_ domain before, during, and after filing the dark spots remains the same and is only affected by the overall photo-bleaching of the dye (see Fig. S7). Thus, the dark spots do not result from the local bleaching of the CTxB label, but rather from the local redistribution/aggregation of the GM1-CTxB complexes.

More detailed insights and the proof for the aggregation of the CTxB within L_o_ domains comes from fluorescence images with the 3-fold labeling. We kept the labeling of the L_d_ and L_o_ phases (DOPE-Atto and GM1-CTxB, respectively) but we added fluorescently labeled cholesterol (TopFluor), which should partition in both L_d_ and L_o_ phases (see Fig. S8). As expected, for domains that exhibit a homogeneous distribution of CTxB within the L_o_ domain, we observe homogeneous colocalization of CTxB and labeled cholesterol within the L_o_ phase. For domains that exhibit aggregation of CTxB, we still observe the homogeneous distribution of the labeled cholesterol. This unambiguously proves that the local appearance of dark areas within the L_o_ phases is solely related to CTxB aggregation and not to structural changes of the membrane. While the exact reason behind the CTxB aggregation remains elusive, it should be noted that it is mainly observed where aggregates of other membrane constituents on top of the membrane are present. We also note that at about 50% RH, aggregates on top of the membrane break into smaller pieces, likely taking up the energetically more favorable structure at the anhydrous conditions. Intriguingly, when increasing the hydration state of the membrane, the homogeneous fluorescence signal within the L_o_ domains is recovered, indicating that the distribution of the GM1-CTxB complexes becomes homogeneous (Fig. S3).

It is evident that the dehydration process itself, when carried out in a controlled manner, does not affect the structure of the SLB. Such preserved membrane structure-wise remains insensitive to dehydration and rehydration cycles.

### Dynamics

With a decrease in hydration level, the mobility of L_d_ lipids decreases. As evident from Fig. 3B, we find that the diffusion coefficient decreases between the fully hydrated sample and the fully dehydrated sample by over a factor of 6 (from 1.69 to 0.27 μm^2^/s), which confirms a major role of water in lipid dynamics.

Upon removal of bulk water when the SLB is equilibrated to a humid environment (90% RH), the diffusion coefficient remains unchanged and the fluorescence intensity recovers to a similar extent after photobleaching, as in the case of fully hydrated SLB (Fig. 3C). This implies that the fluidity of L_d_ lipids remains unhindered in the absence of bulk water and that water molecules present per lipid at 90% RH are sufficient for the lipids to retain their native (read in full hydration) mobility. This is understandable, as at high RH membrane constituents can coordinate as many water molecules as it is energetically most favorable, likely completely filling their direct hydration shell. The biggest changes to the diffusion coefficient are observed with lowering the RH down to about 50%. Further lowering of RH brings little change to the diffusion coefficient. In order to understand this behavior we need to consider the hydration structure of individual lipid molecules.

Phosphatidylcholine (PC) molecules are zwitterionic lipids containing a positively charged choline ((CH_3_)_3_N^+^CH_2_CH_2_OH) moiety and a negatively charged phosphate (PO_4_^3-^) group. Three distinct regions have been identified (Fig. 4) within PC, where water molecules are bound either by weak van der Waals interactions or H-bonds^57^. Region I corresponds to the interior water molecules, buried deeper in the membrane and forming H-bonds with carbonyl oxygens of the glycerol region. Region II refers to water molecules forming a cage-like (clathrate) structure around the whole phosphocholine group. Finally, the consecutive hydration shells/bulk water around the head group belongs to region III^20^. Molecular dynamics (MD) simulations of a PC bilayer revealed that there are 10-12 water molecules in the 1^st^ hydration shell and among these, up to 3 water molecules remain tightly attached to lipids (in the glycerol and/or phosphate region), even after drastic drying^19^. Another MD simulations by Gierula et al. showed that non-esterified oxygen atoms of phosphate group form ~4 H-bonds and the two carbonyl oxygen atoms form ~1 H-bond each^21^, thus ~6 H-bonded water molecules per PC are present in the 1^st^ hydration shell (region I and II). The choline group cannot form a H-bond with water molecules, instead remains surrounded by clathrate hydrate containing ~6.4 water molecules^21^, held via weak electrostatic and van der Waals interactions.

**Figure 4.**
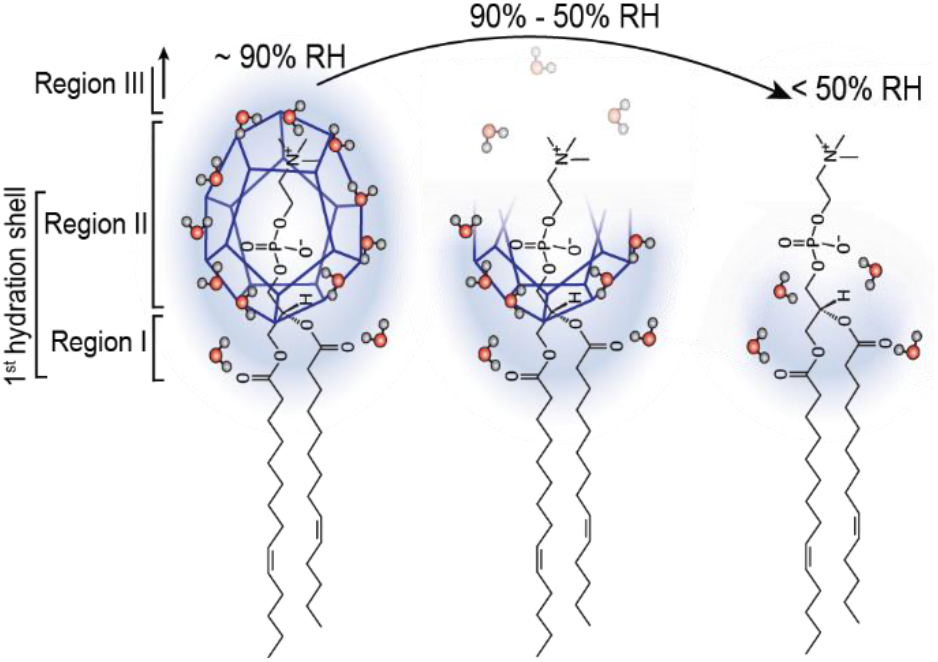
Schematic description of stepwise detachment of water molecules from the three regions of 14:1 PC lipid during controlled dehydration process.

The experimental studies using XRD and infrared spectroscopy also showed that upon bulk water dehydration of stacked lipid bilayers and equilibration of the system at ~95% RH there are around 11-12 water molecules per lipid, confirming quantitatively the structure of the first solvation shell around the lipid group^16,58^. Both, theoretical and experimental studies are thus consistent as to the number of water molecules (~12) per PC lipid in the 1^st^ solvation shell. The same experimental works determined that further decrease in RH (95% → 75% → 50% → 25%) of the environment of bilayers stacks leads to lowering of the hydration of lipids to approximately 10.9, 6.3, 3.6, 2.4 water molecules per lipid (averaged from the two experimental works), respectively. Naturally, the desorption of water molecules should occur according to their H-bonding energies. Previous studies reported that H-bonds between the water molecule and the phosphate group are stronger than water-carbonyl group H-bonds, while both of these H-bonds are stronger than inter-water molecules H-bonds^57^. Therefore, water molecules loosely bound with weak van der Waals interactions, as well as, bound to other water molecules will detach first, followed by a detachment of water molecules bound through strongest H-bonds to phosphate and/or carbonyl moieties.

Supplementing our experimental observations with the considerations above, a clear picture of the interplay between the water and the lipid membrane emerges (Fig. 4). Upon withdrawal of bulk water and equilibration of the SLB with high RH, outer solvation shells are removed and only the first, direct solvation shell containing around 12 water molecules per lipid remains. Under these conditions, the diffusion coefficient of L_d_ phase remains unaffected. Clearly, the water molecules beyond the first hydration shell are not involved in the mobility regulation of the lipids in SLBs. When decreasing the RH down to ~50%, a sharp and continuous drop in the mobility of L_d_ phase lipids occurs. In this regime, each lipid loses 6-7 water molecules. This implies that the clathrate structure breaks apart because at 50% RH only about 4 water molecules are left, which is insufficient to form a stable cage around the phosphate moiety. In the regime below 50% RH the lipid mobility hardly changes. Apparently, the remnant 2-4 water molecules tightly attached to phosphate and, in particular, carbonyl oxygen atoms do not affect lipid mobility. It is evident that out of the water molecules within the 1^st^ solvation shell, those forming the clathrate structure are mostly involved in facilitating the lateral diffusion of the lipids in SLBs.

After establishing which water molecules contribute to the regulation of the lateral mobility of lipids, the question arises why and how these water molecules affect the mobility of lipids. For each diffusion step, a lipid molecule needs to possess energy higher than the activation energy of diffusion (E_a_) and to have sufficient free volume in the vicinity^59^. Free volume in our SLBs could decrease if small perforations (or nano holes) were formed in the bilayer in a dehydrated condition. However, fluorescence images and AFM topography images (Fig. S5), revealing flawless and uniform L_d_ phase in dehydrated SLBs, nullify this scenario. Therefore, the activation energy factor must dominate here.

Water molecules forming the clathrate screen the repulsive Coulombic interactions between adjacent lipid head groups^19,60^. Consequently, in the absence of this shielding water cage, the repulsive interactions between adjacent head groups become more prominent, increasing the activation energy for diffusion. In other words, for dehydrated SLB, a lower population of lipids possesses sufficient energy to overcome the diffusion activation energy barrier. Consequently, the probability for a lipid molecule to overcome the activation energy barrier at a particular time decreases, leading to an overall decrease in diffusion coefficient and mobile fraction. We confirmed this by measuring the activation energy for diffusion for fully hydrated and dehydrated (~30% RH) SLBs. Figure 5A depicts representative Arrhenius plots for hydrated and dehydrated SLB. E_a_ for fully hydrated bilayers averaged over 4 datasets (2 SLBs, increasing and decreasing temperature for each SLB), amounts to 23±4 kJmol^-1^, which is consistent with the previous reports^61^. E_a_ for dehydrated membrane, averaged over 4 datasets, is approximately twice as high and amounts to 47±17 kJmol^-1^. An increase of E_a_ for dehydrated lipid monolayer has been qualitatively predicted earlier based on theoretical considerations^62^. The higher standard deviation of E_a_ for dehydrated sample results from higher uncertainty in fitting the very slow fluorescence recovery in the FRAP data. Evidently, decreasing hydration of the SLB leads to a noticeable increase in activation energy for the L_d_ phase.

**Figure 5.**
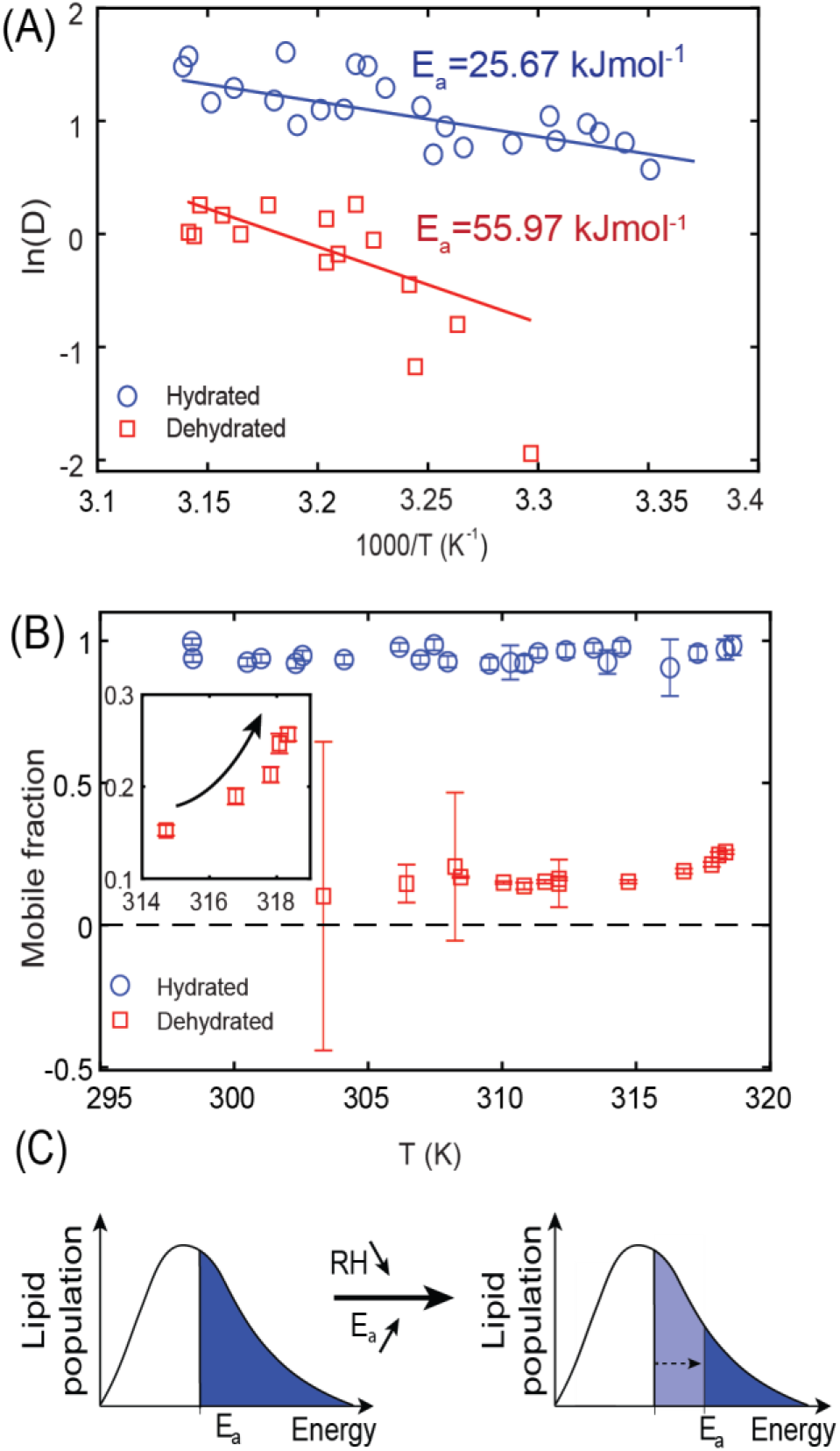
(A) Arrhenius plots for one representative fully hydrated (blue circles) and dehydrated (red squares) SLB. (B) Temperature dependence of the mobile fraction extracted from the data shown in panel A. (C) Schematic representation of the relation between the diffusion activation energy and the lipid mobile fraction in the dehydration experiment.

Importantly, the significant increase in E_a_ for diffusion with dehydration explains the observed decrease in mobile fraction during dehydration. As E_a_ increases, the population of molecules having enough energy to overcome the barrier at a particular time decreases (Fig. 5C), which is reflected in the slower recovery of FRAP traces and lower mobile fraction. In this case, an increase in mobile fraction should be observed with increasing the temperature, as more energy is delivered to the lipids. For a fully hydrated sample, the mobile fraction is already >~95% and there is very little or no room for it to increase further. In the case of dehydrated SLB, the mobile fraction indeed tends to increase at higher temperatures (Fig. 5B).

Altogether, the observed slowing down of the diffusion with an increase in diffusion activation energy suggests that the SLB quasi-gellifies (stiffens) upon dehydration, in particular upon the removal of the clathrate water molecules. This is in agreement with previous studies, which indeed suggested that dehydration leads to an increase in the main phase transition temperature of lipids^63,64^ indicating fluid-to-gel like transition at lower hydration conditions.

Finally, we note that with rehydration, the former mobility of lipids is restored. The variation of diffusion coefficient with hydration state for few dehydration and rehydration cycles demonstrates that the mobility of lipids strongly depends on the availability of the water molecules per lipid, and the diffusion coefficient is instantaneously responsive towards water content. It is also evident that for the intact membrane that survived the dehydration process, losing or gaining mobility of lipids due to change in hydration is completely reversible and repeatable.

## Conclusions

We successfully prepared desiccation-tolerant, phase separated lipid bilayers without mechanical or chemical alterations. While a rapid reduction in water content causes irreversible damage to the SLB structure, a gradual and controlled dehydration process allows the preparation of stable SLBs even in the complete absence of water. Dried SLBs can be brought back to full hydration without affecting their integrity and reused as functional membranes after rehydration. Thus, storage and handling of such desiccation-tolerant SLBs become much easier for bioengineering applications such as biocoatings.

We carefully addressed the structural and dynamical properties of SLBs across a wide spectrum of hydration states. While structurally, SLBs showed little sensitivity to the hydration state of the SLB, we observed a 6-fold decrease in lateral diffusion coefficient for the lipids forming L_d_ phase with lowering hydration of the SLB. Importantly, we correlated the observed changes of the diffusion coefficient with the lipid hydration structure and established that these are 6-7 water molecules hydrating the phosphocholine headgroup and forming a cage-like structure that act as a *lubricant* for the diffusion and modulate the lateral mobility of disordered phase in the SLBs. We demonstrate that the observed slowing down of the diffusion is directly related to an increase in activation energy for diffusion at lowered hydration conditions. Together with the unpredicted overall structural stability, these findings point towards quasi-gellification of the SLB with lowering its hydration. Intriguingly, the native dynamics of fully hydrated SLB is recovered with rehydration. Consequently, the dried SLB with unperturbed membrane structure and dramatically reduced mobility can be considered as a less active form of the membrane, which can be compared with the dormant stage of organisms exhibiting anhydrobiosis. Local ‘anhydrobiosis’ occurs also in our organisms during for instance cell-cell interactions or during binding of large biomacromolecules, when the water molecules are expelled from the interaction site. It is thus conceivable that the observed slowing down in SLB dynamics also occurs locally and leads to stiffening and stabilization of the membrane, potentially stabilizing transiently molecular interactions.

Our studies on the interplay between the membrane and its hydration open up a range of exciting experiments that could certainly provide new molecular-level insights into effects such as hydrophobic mismatch, line tension, or the properties of the inter-phase boundaries of the membrane structural complexes.

Finally, a clear relation between the diffusion coefficient and the number of water molecules hydrating the membrane could be utilized for biosensing applications to monitor the local hydration state of biomimetic systems. This idea further gains in impact when the hydration sensing is done on a single molecule/probe level using e.g. fluorescence correlation spectroscopy or single particle tracking techniques. Our findings thus hold a huge application potential from both biological and technological viewpoint.

## ASSOCIATED CONTENT

Supporting information: Supplementary Movie 1 and Supplementary Figures S1 – S8.

## AUTHOR INFORMATION

### Notes

The authors declare no competing financial interest.

## ACKNOWLEDGMENT

The authors acknowledge the financial support from the EMBO Installation Grant 2019 and from the Ministry of Science and Higher Education of Poland in the year 2021 under Project No 0512/SBAD/2120. LP acknowledges the financial support from the First TEAM grant No. POIR.04.04.00-00-5D32/18-00, provided by the Foundation for Polish Science (FNP). This work was financed from the budget funds allocated for science in the years 2019–2023 as a research project under the “Diamond Grant” program (Decision No. 0042/DIA/2019/48). The authors thank MSc. Marek Weiss and Prof. Arkadiusz Ptak for their assistance in acquiring AFM images. PS and HGF acknowledge the financial support from the Max Planck Society.

