## Supplementary Figures for "Hydration layer of only few molecules controls lipid mobility in biomimetic membranes"

**Table of contents:**

Supplementary figures S1 – S8.

**Figure S1**

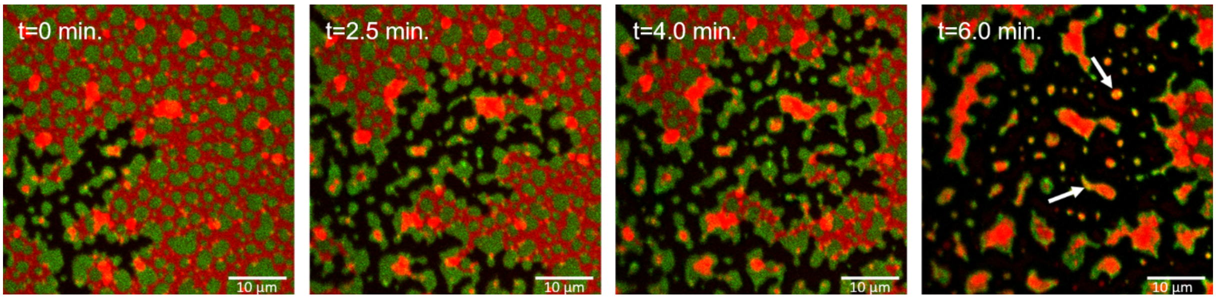


Fig. S1. Consecutive fluorescence images of the same area of SLB exposed abruptly to ambient RH. Desorption of water causes shrinking of the remnant water layer and induces delamination of the lipids from the mica substrate. During drying, the water layer wavefront passes over the surface and causes detachment of the $L_d$ phase. $L_o$ phase, which has a stronger affinity to the mica substrate, stays unperturbed for a longer time. With time both phases pill off the solid support. Dried SLB (rightmost image) contains aggregates composed of both phases (yellow spots, marked with white arrows).

**Figure S2**

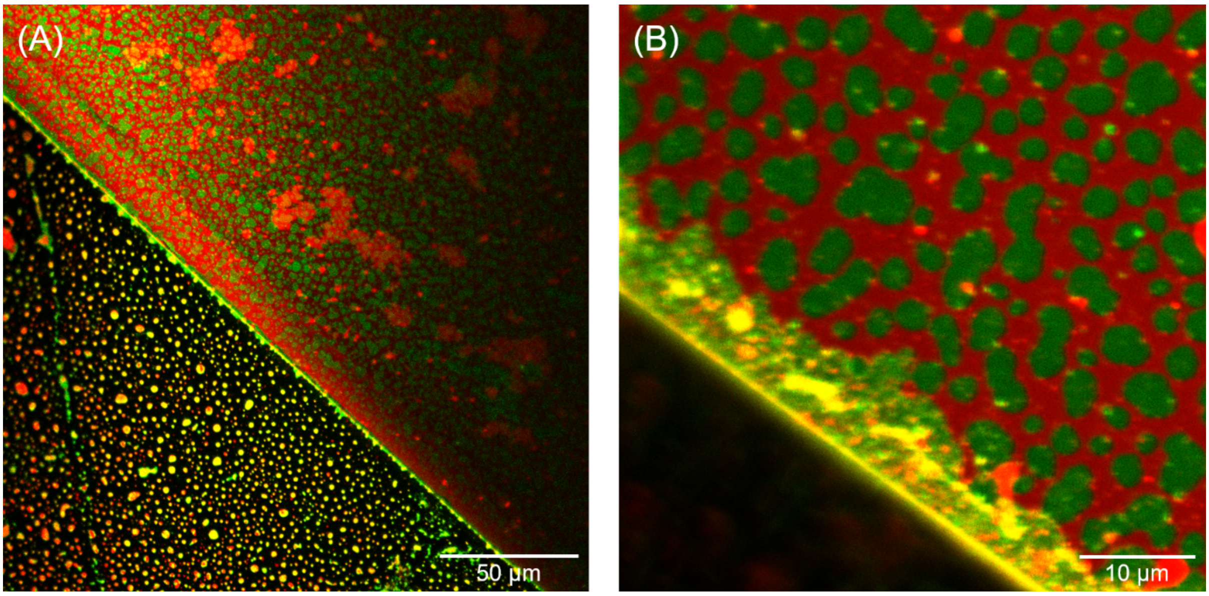


Fig. S2. The presence of defects of the solid support (mica cleaving-induced defects) can influence membrane stability upon dehydration. Mica terraces stop the drying waterfront and act as an obstruction preventing the membrane from delamination. (A) Membrane curls up in the bottom-left part of the image, forming vesicles and aggregates, and remains unperturbed in the upper-right part section of the image, (B) Close-up image of the border between the two regions. During dehydration the dragged lipids were deposited by the moving waterfront along the defect.

**Figure S3**

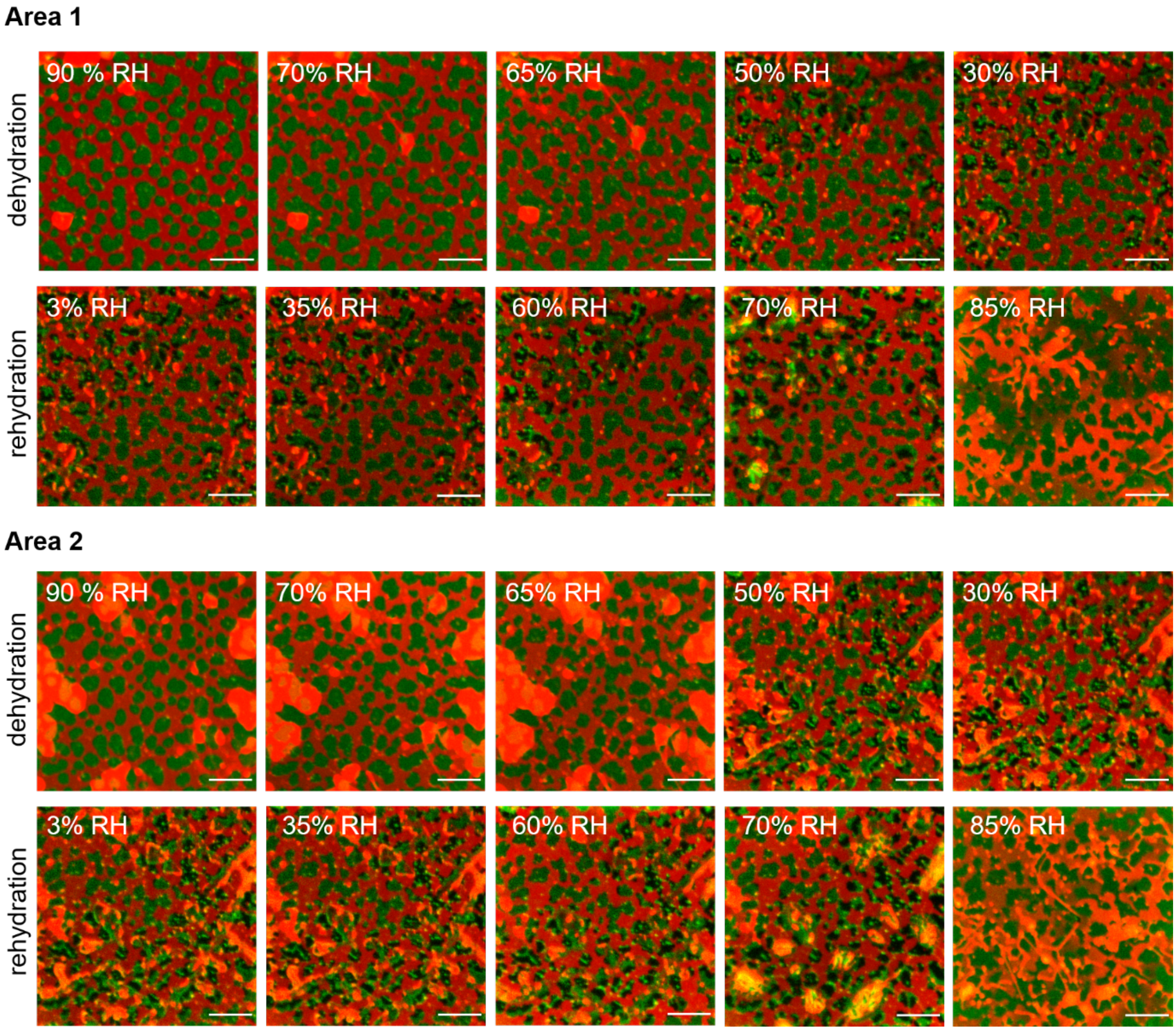


Fig. S3. Consecutive fluorescence images of two different areas (area 1 and area 2) of SLB during dehydration (top row) and rehydration (bottom row). At the humidity of around 50% RH and lower, in some areas of the membrane, local aggregation of the GM1-CTxB occurs, which is visible as the formation of darker and brighter spots within the $L_o$ domains. At about 50% RH aggregates accumulated on top of the membrane break into smaller ones. During rehydration, $L_o$ domains regain homogeneous CTxB distribution (and hence homogeneous fluorescence signal) at about 85% RH. The scale bar is 10 µm.

### Figure S4

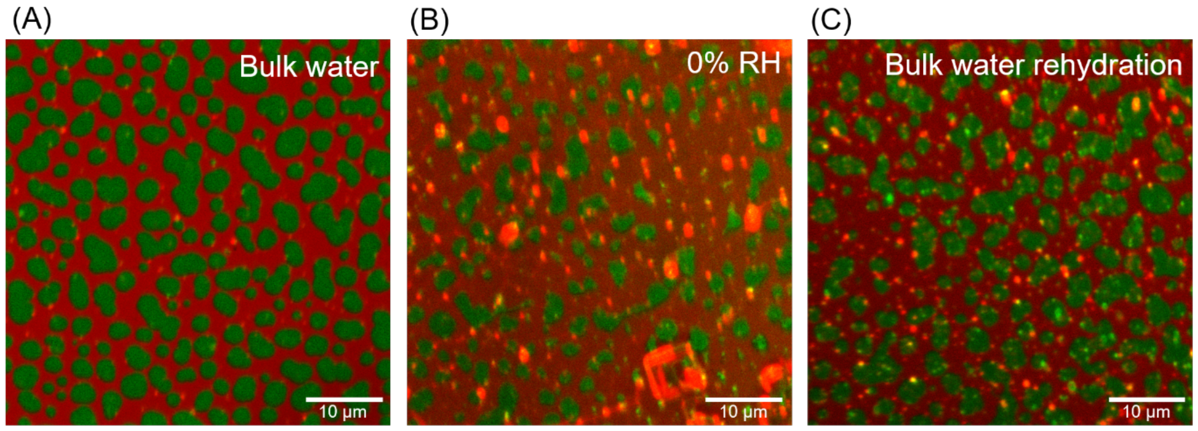


Fig. S4. Confocal images of the same sample recorded at different hydration conditions: (A) right after preparation, full hydration with bulk water, (B) slowly dehydrated by decreasing the humidity level and kept at 0% RH, (C) slowly rehydrated and refilled with bulk water. Membrane structure remains unaffected after a complete cycle of de(re)hydration.

### Figure S5

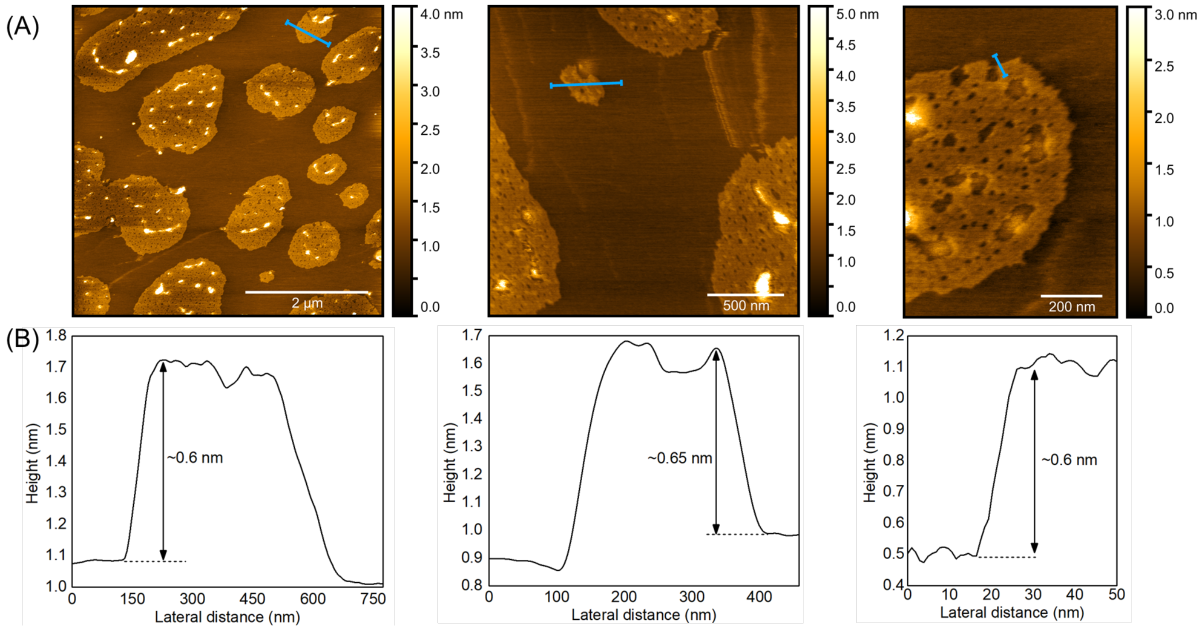


Fig. S5. AFM images of dehydrated SLB. (A) Topography scans confirming intact membrane with continuous $L_d$ phase at dehydrated condition. Presence of nanoscopic depressions with a depth of ~0.5-0.7 nm is noticed in $L_o$ phase domains. Given the same height difference as between the $L_o$ and $L_d$ phases, these depressions correspond to $L_d$ nanodomains trapped in the $L_o$ phase. A similar effect was observed for ceramide-rich membranes, where due to lower line tension of the $L_o$ phase, $L_d$ nanodomains participated in the $L_o$ domains[1]. Elevated features with the height of ~6 nm (from the average surface height of $L_o$ domains) correspond to the GM1-CTxB complexes. Scale bars are 2 μm, 0.5 μm and 0.2 μm for the left, middle and rightmost image, respectively. (B) Cross-sections profiles highlighting the height difference between the $L_d$ and $L_o$ phase (marked by blue lines in panel A).

**Figure S6**

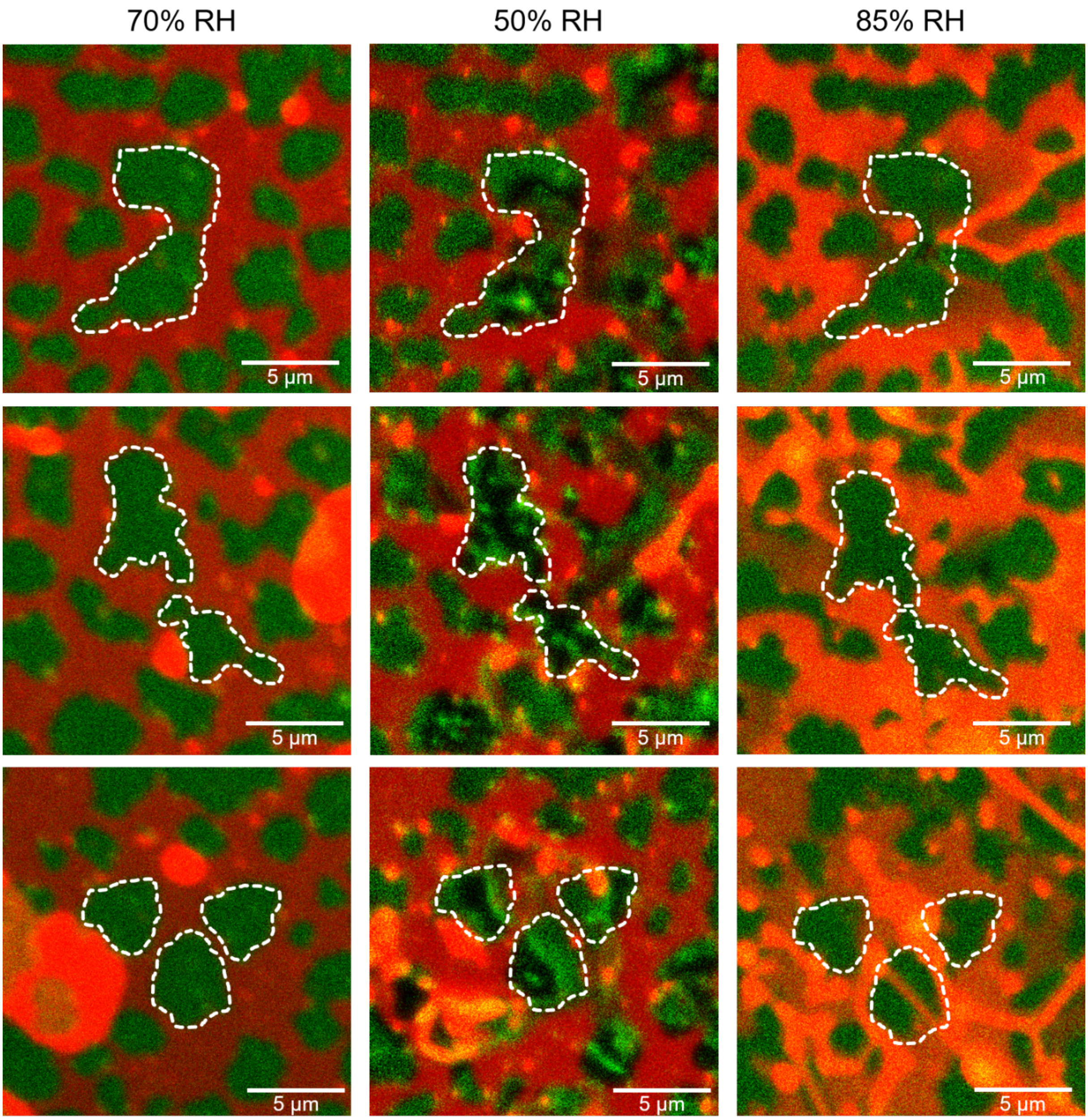


Fig. S6. SLB consisting of $L_d$ and $L_o$ phase exposed to the relative humidity of 70% and 50% (during dehydration) and 85% (during rehydration). The white, dashed lines mark the outlines of the domains. The shape and area of the domains during the dehydration and rehydration processes remain largely the same. The GM1-CTxB complexes aggregate at around 50% RH but upon rehydration redistribute again evenly within the whole domain.

**Figure S7**

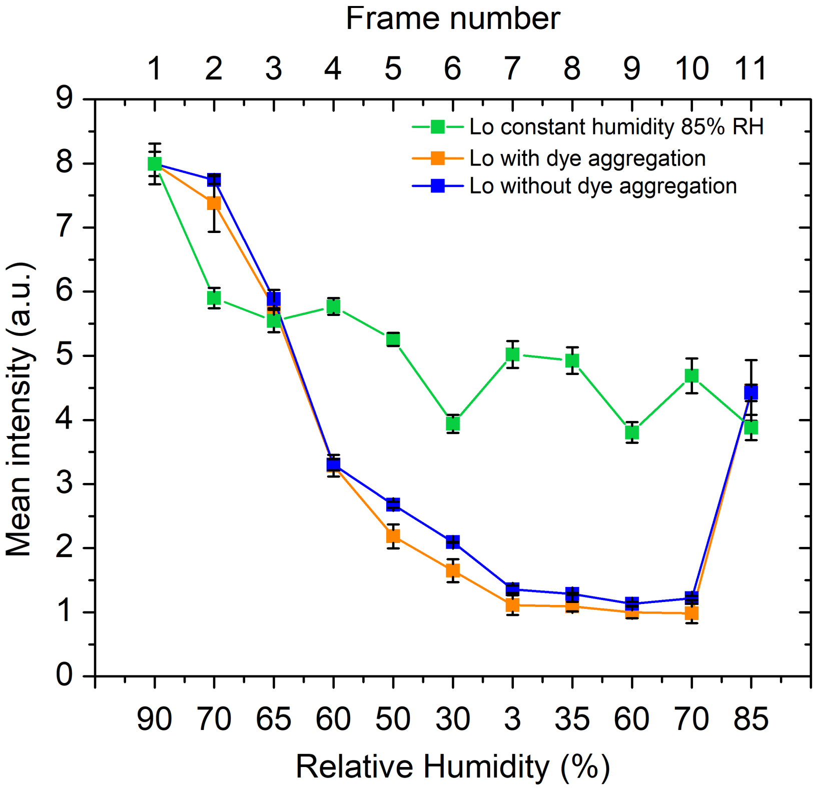


Fig. S7. Changes in the mean intensity of the $L_o$ domains with de(re)hydration. Green trace reflects the native fluorescence bleaching of the Alexa Fluor 488 dye caused by the consecutive imaging of the SLB exposed to the constant humidity of 85% RH. The mean intensity is plotted as a function of frame number (top x-scale). Orange trace shows the changes in the fluorescence intensity of the domain showing aggregation of the GM1-CTxB complexes during dehydration and rehydration. Blue trace represents the changes in the domain intensity for a domain that did not show any aggregation of the GM1-CTxB complex. The overall fluorescence intensity for the domains with and without the aggregation changes in the same manner but it shows a more rapid intensity decrease than for the sample kept in constant humidity. Upon rehydration, at 75-85% RH, the fluorescence of the domain is restored. The final mean intensity of the fluorescence after taking 11 images at constant humidity is the same as after the whole de-/rehydration cycle. Evidently, the strong decrease in fluorescence of the Alexa Fluor 488 dye is due to the lower quantum efficiency of the dye at lower hydration and not solely due to photobleaching of the dye. Lipid domains were analyzed using ImageJ software by measuring the mean intensity from the chosen $L_o$ domains (three showing GM1-CTxB aggregation and three with homogeneous GM1-CTxB distribution). Intensities were normalized with respect to their initial value.

**Figure S8**

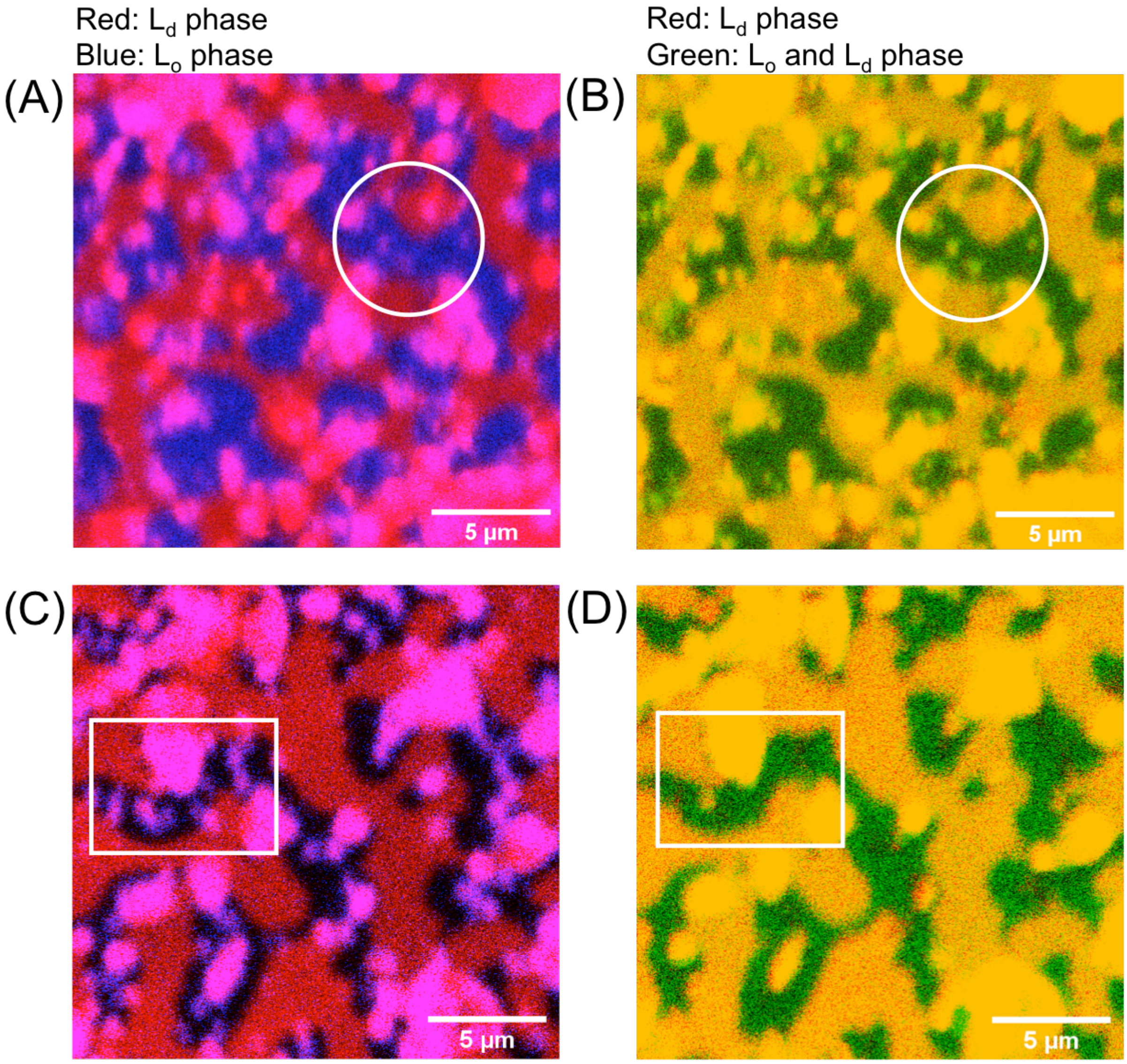


Fig. S8. Confocal images of two areas of SLB equilibrated at low relative humidity (~30% RH), where $L_o$ domains show homogeneous (top row, A-B) and inhomogeneous (bottom row, C-D) distribution of the GM1-CTxB. (A-C) $L_d$ phase is labeled with Atto-633-DOPE (red) and $L_o$ phase is labeled with CTxB-Alexa594 (blue). (B-D) $L_d$ phase is labeled with Atto-633-DOPE (red), while cholesterol, which partitions in both $L_d$ and $L_o$ phase is labeled with the TopFluor dye (green). $L_d$ phase appear yellow in panels B-D, as it overlaps homogeneously with the green signal assigned to the cholesterol label. It is clear that, while in some areas GM1-CTxB tends to aggregate leaving dark (no fluorescence) spots within the $L_o$ domains (panel C, white rectangle), the structure of the $L_o$ domains themselves remains intact. This is evident from the unperturbed, homogeneous distribution of the cholesterol-TopFluar (panels B-D).
